# Comparison of TRIBE and STAMP for identifying targets of RNA binding proteins in human and *Drosophila* cells

**DOI:** 10.1101/2023.02.03.527025

**Authors:** Katharine C. Abruzzi, Corrie Ratner, Michael Rosbash

## Abstract

RNA binding proteins (RBPs) perform a myriad of functions and are implicated in numerous neurological diseases. To identify the targets of RBPs in small numbers of cells, we developed TRIBE, in which the catalytic domain of the RNA editing enzyme ADAR (ADARcd) is fused to a RBP. When the RBP binds to a mRNA, ADAR catalyzes A to G modifications in the target mRNA that can be easily identified in standard RNA-sequencing. In STAMP, the concept is the same except the ADARcd is replaced by the RNA editing enzyme APOBEC. Here we compared the two enzymes fused to the RBP TDP-43 in human cells. Although they both identified TDP-43 target mRNAs, combining the two methods more successfully identified high confidence targets. We also assayed the two enzymes in *Drosophila* cells: RBP-APOBEC fusions generated only low numbers of editing sites, comparable to the level of control editing. This was true for two different RBPs, Hrp48 and Thor (*Drosophila* EIF4E-BP), indicating that TRIBE performed better in *Drosophila*.

## INTRODUCTION

RNAs are bound by RNA binding proteins (RBPs) even in the nucleus before transcription is complete. For example, the association of RBPs with pre-mRNA affects all aspects of nuclear RNA processing, from capping, splicing and 3’ end formation to nuclear export. Additional RBPs bind to cytoplasmic mRNAs and determine their subcellular localization and stability. Yet other RBPs act to regulate translation including the association of specific mRNAs with the ribosome. The coordinated functioning of these RBPs is necessary to generate the right amount of the correct protein, at the right time and in the right place.

Many human neurological diseases are linked to RBP mutations. This vulnerability may be due in part to the fact that most central brain neurons are long-lived and cannot divide themselves out of trouble. For example, mutations in FMRP are responsible for Fragile X syndrome (Penagarikano et al. 2007), dysregulation of TDP-43 underlies amyotrophic lateral sclerosis (ALS, Ling et al. 2013; Gao et al. 2018), and mutations in SMN1 cause spinal muscular atrophy (SMA, Farrar and Kiernan 2015). It is therefore important to understand the mRNA targets of key neuronal RBPs.

For many years, Crosslinking Immunoprecipitation (CLIP) has been the tried-and-true method for identifying the targets of RNA binding proteins. CLIP is a powerful method because UV crosslinking is used to biochemically attach the RBP to the target mRNA, which makes precise binding site identification possible. However, CLIP also has drawbacks: an efficient antibody is needed, crosslinking is biased to guanosine and thymidine residues, crosslinking can capture weak or transient interactions, and a very large amount of biological material is usually required (reviewed in Hafner et al. 2021; Xu et al. 2022). As we began to understand more about the heterogeneity of tissues and even single cells, it became apparent that a method that can identify RBP targets in small amounts of material was needed.

In the past seven years, two methods have been developed that allow researchers to examine the targets of RBP in small discrete groups of cells. The first was from our lab in 2016 and is called TRIBE (<u>t</u>argets of <u>R</u>NA binding proteins *i*dentified <u>b</u>y <u>e</u>diting; McMahon et al. 2016). In TRIBE, an RBP of interest is fused to the catalytic domain of ADAR (ADARcd). When the RBP binds its target mRNA, the ADARcd edits the mRNA. In 2018 we incorporated a single point mutation in the ADARcd, which substantially increased editing activity and decreased local sequence preferences (Kuttan and Bass 2012); this variant has been used for all recent TRIBE experiments (HyperTRIBE; Xu et al. 2018). Editing sites are then identified computationally from RNA sequencing data. Importantly, this allows target identification in very small numbers of cells and neurons without requiring biochemistry. TRIBE has been useful to a number of other labs working in different fields and model organisms ranging from humans to malaria parasites (Liu et al. 2019; Alizzi et al. 2020; Nguyen et al. 2020; Arribas-Hernández et al. 2021; Cheng et al. 2021; Singh et al. 2021; van Leeuwen et al. 2022).

A very similar strategy, conceptually and in overall design, was recently published in which a RBP is fused to a different editing enzyme, APOBEC1 (apolipoprotein B mRNA editing enzyme, catalytic polypeptide-like). This method was used in two different studies and named STAMP (Brannan *et al*., 2021) and Dart-seq (Meyer, 2019); we will use the nomenclature STAMP in this paper. TRIBE and STAMP both offer distinct advantages and disadvantages. ADAR is an adenosine deaminase and makes A-to-I edits within a supposedly double-stranded region even without the RNA binding regions normally present in ADAR (Macbeth et al. 2005). APOBEC1 is a cytidine deaminase and therefore catalyzes C-to-T edits in single-stranded RNA but also edits single-stranded DNA (Rosenberg et al. 2011; Smith et al. 2012; Salter et al. 2016). ADAR and APOBEC both have some local sequence preferences (U<u>A</u>G ->ADAR; A/U flanked ->APOBEC) (Rosenberg et al. 2011; Kuttan and Bass 2012). TRIBE only fuses the ADARcd, which can be cleanly separated from the ADAR double-stranded RNA binding regions to the RBP. The latter normally directs the specificity of full-length ADAR and is replaced by the RBP in TRIBE. In contrast, the entire APOBEC protein is fused in STAMP, perhaps because it is uncertain how the editing specificity of APOBEC is defined. For example, APOBEC-mediated editing may require dimerization as well as cofactors (e.g., A1CF and RBM47; reviewed in Smith et al. 2012; Fossat et al. 2014).

To compare the ability of TRIBE and STAMP to identify targets of RBPs, we examined these methods side-by-side in HEK-293 and *Drosophila* cells. In HEK cells, the human ADAR2cd (referred to in this manuscript as ADAR) and rat APOBEC1 (referred to in this manuscript as APOBEC) were fused to Tar DNA binding protein-43 (TDP-43). We previously successfully used this protein for TRIBE in a mammalian system (Herzog et al. 2020).

As indicated by published STAMP results (Brannan et al. 2020), APOBEC worked well in HEK cells. TDP-43-APOBEC generated a similar number of editing sites as TDP-43-ADAR, greater than 10-fold more sites than expressing the editing enzymes alone. TDP-43-ADAR and TDP-43-APOBEC both identified substantial numbers of target genes, although TDP-43-ADAR identified 50% more genes than TDP-43-APOBEC. Moreover, 70% of the TDP-43-APOBEC target transcripts were also identified by TDP-43-ADAR. We also compared STAMP and TRIBE in *Drosophila*, with two RBPs, Hrp48 and Thor (*Drosophila* EIF4E-BP). Their targets were both successfully identified with TRIBE (McMahon et al. 2016; Jin et al. 2020). To test STAMP, we replaced the ADARcd with APOBEC to generate Hrp48 and Thor STAMP. Although TRIBE worked well with both of these RBPs as expected there was no substantial editing of target mRNAs with STAMP.

## RESULTS

To directly compare the efficacy of ADAR (TRIBE) with that of APOBEC (STAMP) in HEK-293 cells, we took advantage of two existing TDP-43-ADAR TRIBE constructs (Herzog et al. 2020): the CMV promoter expressed either the TDP-43 coding sequence followed by the ADARcd or the ADARcd alone (pCMV-TDP-43-ADARcd and pCMV-ADARcd). Parallel constructs were generated in which the ADARcd was replaced with rat APOBEC1 as used in STAMP experiments (Fig. 1A, Brannan et al. 2020; pCMV-TDP-43-APOBEC and pCMV-APOBEC). These plasmids were transfected into HEK-293 cells along with a pCMV-eGFP plasmid. As a control, HEK cells were transformed with pCMV-EGFP only. Twenty-four hours after transfection, GFP-positive transfected cells were isolated using a Melody Fluorescence Activated Cell sorter (BD Melody FACS). RNA sequencing libraries were generated from two different biological replicates using Smart-seq2 (see Materials and Methods).

**Figure 1.**
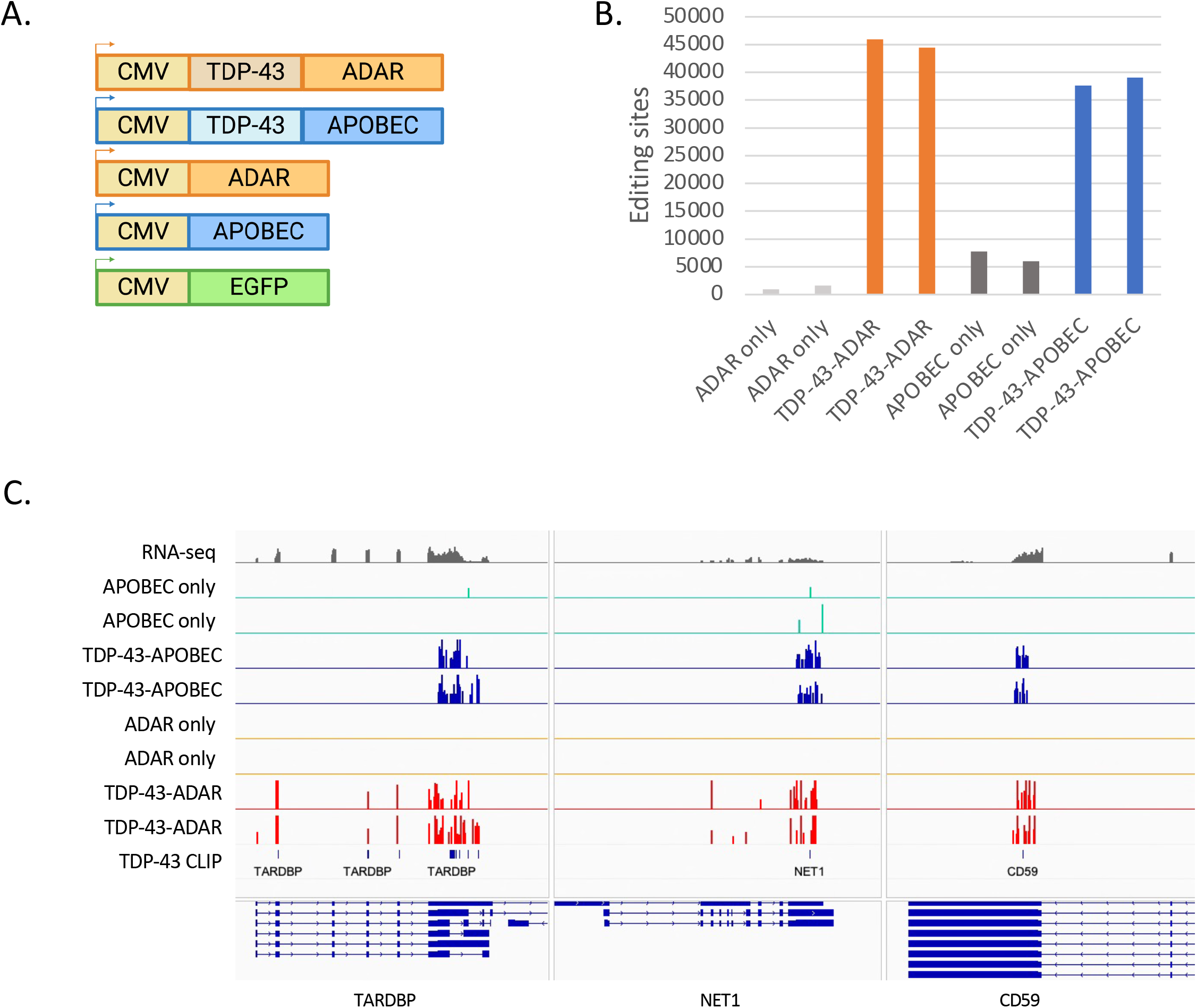
Both TDP-43 TRIBE (ADAR) and TDP-43 STAMP (APOBEC) identify candidate TDP-43 targets in HEK-293 cells. A. Design of constructs for the expression of TDP-43-ADAR, TDP-43-APOBEC and their respective controls in HEK-293 cells. All transgenes were expressed using the Cytomegalovirus promoter (CMV). B. Expression of TDP-43-ADAR and TDP-43-APOBEC in HEK-293 cells generated RNA editing sites above the background level identified by either ADAR or APOBEC alone. The graph quantifies A to G transitions for ADAR and C to T transitions for APOBEC. Two biological replicates are shown. C. Visualization of RNA editing sites generated by TDP-43-ADAR (red) and TDP-43-APOBEC (blue) on TARDBP (the gene encoding TDP-43), NET1 and CD59. The integrated genomics viewer (IGV; Robinson et al. 2011; Thorvaldsdottir et al. 2013) visualization of two biological replicates of TDP-43-ADAR (red), TDP-43-APOBEC (blue), ADAR alone (orange), and APOBEC alone (green). RNA sequencing data is shown on top in gray. TDP-43-CLIP sites are shown in the bottom track (Hallegger et al. 2021). The direction of transcription is denoted by the arrows on the refseq genes. Exons are shown as closed blocks and introns as lines.

Existing TRIBE computational pipelines were used to identify both A to G editing events (TDP-43-ADAR and ADAR only) as well as C to T editing events (TDP-43-APOBEC and APOBEC only) in the RNA sequencing data (McMahon et al. 2016; Rahman et al. 2018; see Materials and Methods). In short, libraries were trimmed and mapped to the human genome (GRChg38.p13). Differential gene expression analysis was performed to verify that expression of TDP-43-ADAR and TDP-43-APOBEC did not cause dramatic changes to the transcriptomes of the HEK cells (Sup. Fig. 1). To identify editing sites, the numbers of A, G, C and T at every position in the genome were uploaded to a mysql database. To be considered an editing site, a particular genomic location needed to be encoded by predominantly the non-edited base (adenosine (ADAR) or cytosine (APOBEC)) in the HEK cells expressing only EGFP (see Materials and Methods). If this criterion was met, the same genomic location was examined in the experimental samples. To score as edited, a location required at least 20 reads and greater than 6% A to G editing (ADAR) or C to T editing (APOBEC).

Expression of TDP-43-ADAR and TDP-43-APOBEC in HEK cells resulted in 35000-45000 editing sites in both biological replicates of the sequencing libraries (Fig. 1B). The TDP-43-editing enzyme fusions generated substantially more editing than expressing the editing enzymes alone (in grey), showing that RNA editing events are substantially increased by fusing these enzymes to an RBP. As a preliminary investigation of editing site location, we examined these sites on three TDP-43 target transcripts: a known target of TDP-43 (TDP-43 mRNA itself; the gene encoding this mRNA is TARDBP; Ayala et al. 2011; Polymenidou et al. 2011) as well as two novel targets, Net1 and CD59 (Fig. 1C). TDP-43-ADAR (red; two biological replicates) and TDP-43-APOBEC (blue; two biological replicates) edit similar mRNA regions of all three genes. These editing clusters are often co-localized with putative TDP-43 binding sites discovered by CLIP (bottom panel Fig.1C; lines indicate CLIP target sites from Hallegger et al. 2021). Expression of the editing enzymes alone (APOBEC, green; ADAR, orange; two biological replicates each) resulted in many fewer editing sites on these target mRNAs.

Although the number of RNA editing sites generated by TDP-43-ADAR and TDP-43-APOBEC were quite similar, the two enzymes show different editing characteristics. First, expressing APOBEC alone generated 5-fold higher levels of enzyme only editing than expressing ADARcd alone (Fig. 1B). This may be because the entire APOBEC coding sequence was used compared to only the catalytic domain of ADAR (See Discussion.) Second, ADAR more often edits the exact same nucleotides in two biological replicates (48% of sites are identical; another 17% are within 100bp), whereas APOBEC is more likely to edit nearby nucleotides (only 31% of sites are identical and 37% are within 100bp; compare Fig. 2A and 2B). Third, ADAR generates an overall higher level of editing at each site (Fig. 2C). The average percentage editing of a TDP-43-ADAR site is 15.5% compared to 9.8% for TDP-43-APOBEC. This is undoubtedly related to the observation that ADAR often edits the same nucleotide in replicate experiments, i.e., RBP-ADAR is more likely to edit the same A in another copy of the same transcript rather than nearby locations, which results in more reproducibility and a higher editing percentage. Fourth, TDP-43-ADAR yields fewer edited nucleotides per mRNA (mean of 4.8) compared to TDP-43-APOBEC (mean of 7; Fig. 2D). This observation is also because TDP-43-ADAR is more likely to edit the same nucleotide, which yields higher editing percentages at this location but fewer overall edited nucleotides on a target mRNA.

**Figure 2.**
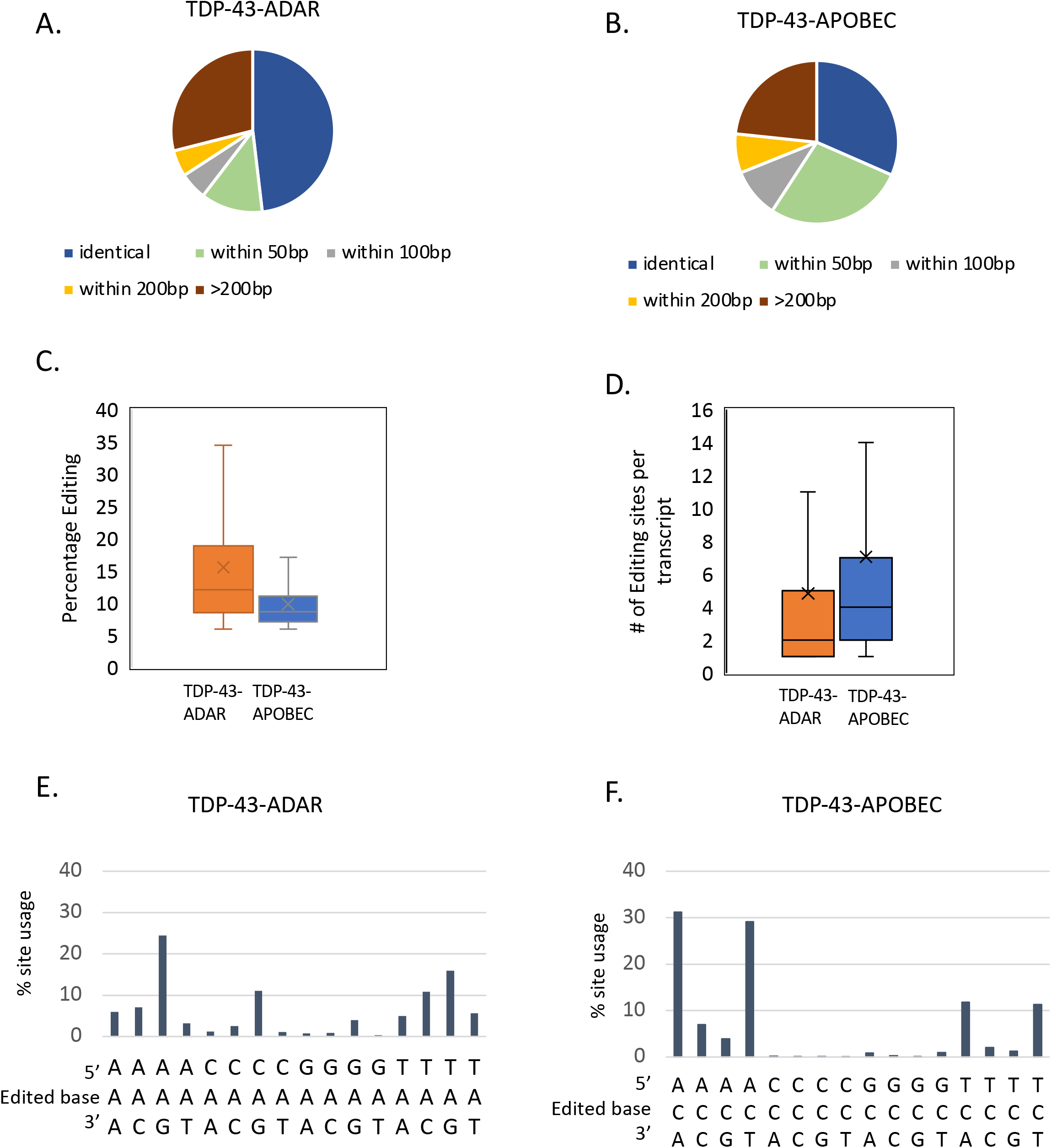
TDP-43 TRIBE and TDP-43 STAMP identify editing sites with different characteristics on TDP-43 target transcripts. A and B. When two biological replicates were compared, TDP-43-ADAR was more likely to edit identical locations in both experiments (dark blue) while TDP-43-APOBEC was more likely to edit two different nucleotides that are in close proximity (green and gray; 50-100 bp). C. TDP-43-ADAR editing sites had a higher mean percentage editing (~15%) compared to those of TDP-43-APOBEC (mean editing ~10%). Mean values indicated by X and median values by lines (p-value <0.0001; Welch’s t-test). D. TDP-43-APOBEC generated ~7 editing sites per transcript (mean value) and TDP-43-ADAR generated ~5 editing sites per transcript (mean value; mean indicated by X and median indicated by a line; p-value <0.0001; Welch’s t-test). E and F. TDP-43-APOBEC and TDP-43-ADAR both showed near neighbor preferences for editing site choice. TDP-43-ADAR (E) preferentially edited adenosines followed by guanosine. TDP-43-APOBEC (F) preferentially edited cytosines that were flanked by A or T. 83% of the edited sites generated by TDP-43-APOBEC were ACA, ACT, TCA or TCT. TDP-43-ADAR showed less bias in editing site choice than TDP-43-APOBEC; the top four triplets edited account for 62% of all edited sites.

We hypothesized that this more localized preference of ADAR may be because it has more restricted site choice rules than APOBEC. To compare these rules, we computationally extracted the neighboring nucleotide 5’-and 3’ of the edited sites (Fig. 2E and F). Surprisingly, TDP-43-APOBEC had a more restrictive site choice than TDP-43-ADAR (Fig. 2E and 2F). 84% of TDP-43-APOBEC editing events occurred on a C flanked by only A or T (Fig. 2F; ACA, ACT, TCT, TCA). Editing sites were rarely observed on cytosines with a 5’C or 5’-G. TDP-43-ADAR also showed specific site choice but to a lesser extent: 64% of the TDP-43-ADAR editing sites were concentrated on AAG, CAG, TAC, TAG (Fig. 2E). These observations are consistent with previous work showing that the ADAR2-E488Q catalytic domain prefers T or A 5’-of the editing sites (Kuttan and Bass 2012). The higher restrictive site choice of APOBEC suggests that the localized preferences of ADAR may be due to RNA secondary structure playing a not insignificant role in the site choice of TDP-43-ADAR (see Discussion).

To accommodate these differences in editing between TDP-43-ADAR and TDP-43-APOBEC and to make a pipeline useful for both enzymes, we made several modifications to our previous TRIBE bioinformatics pipeline. To account for the lower reproducibility of the APOBEC editing sites, we required an APOBEC editing site to be within 100bp of a second site in the biological replicate instead of requiring the same site to be identified in two biological replicates as done for ADAR. In addition, we ignored any nucleotide edited at a frequency above 1% in either replicate of the enzyme-only samples. This conservative filtering step accommodates the higher background editing by the APOBEC-only construct and has been previously used in mammalian cells to ensure that editing sites identified with fusion proteins are not false positives (Biswas et al. 2020). After incorporating these two additional filtering steps, a final set of editing sites and target mRNAs were identified for TDP-43-ADAR and TDP-43-APOBEC (Fig. 3A). Although a nearly identical number of editing sites were identified for the two fusion proteins, TDP-43-APOBEC generated more sites per mRNA and therefore fewer target transcripts (Fig. 3B).

**Figure 3.**
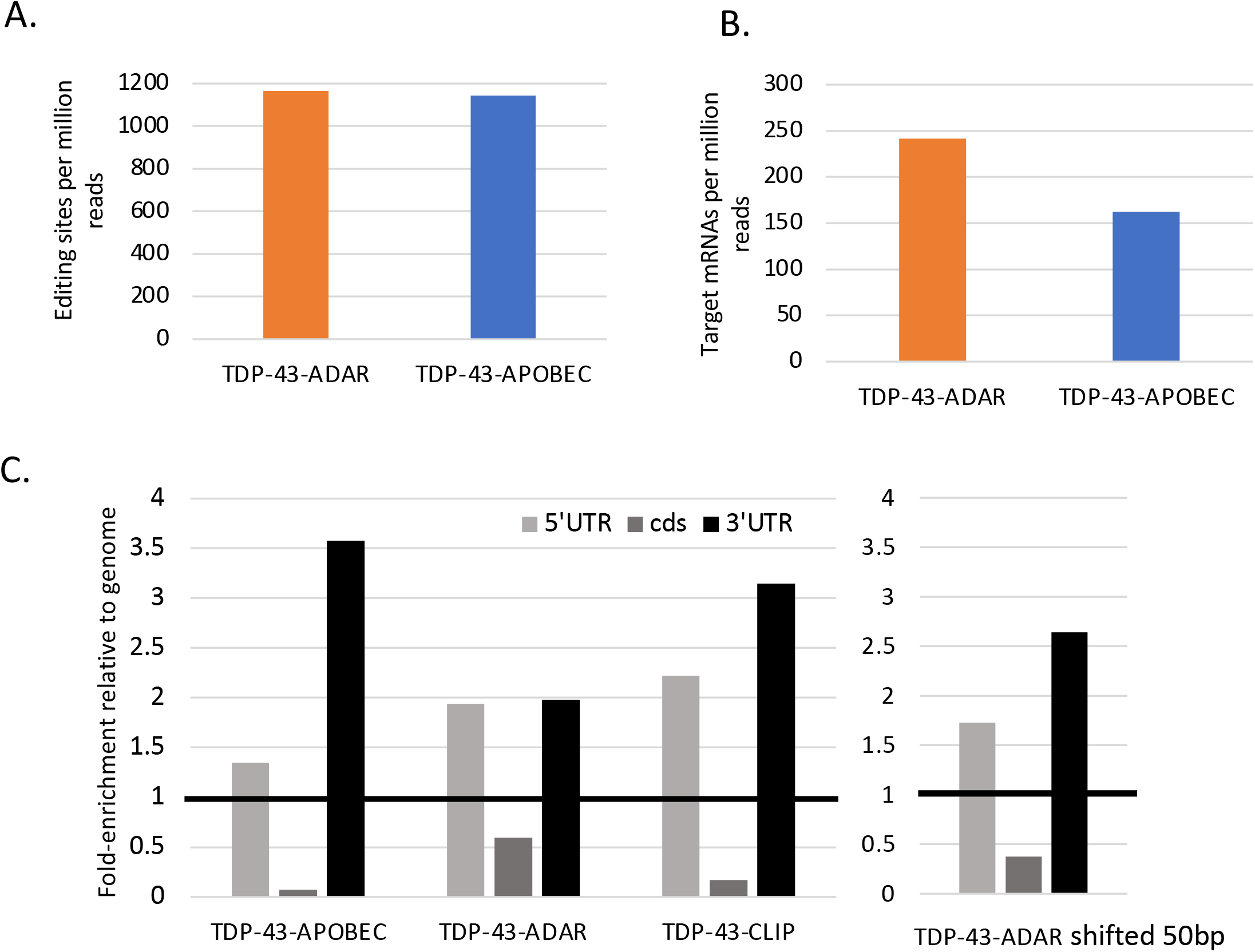
TDP-43-ADAR and TDP-43-APOBEC generated similar numbers of editing sites that were preferentially located in the 3’-UTR and 5’-UTR. A. TDP-43-ADAR and TDP-43-APOBEC generated similar numbers of editing sites. (A to G transitions are indicated for ADAR and C to T transitions for APOBEC). This graph shows only editing sites that were consistent between two biological replicates not found in enzyme only controls. The number of editing sites identified was normalized per million reads to adjust for sequencing library depth. B. TDP-43-ADAR identified more target transcripts than TDP-43-APOBEC. C and D. TDP-43-ADAR and TDP-43-APOBEC editing sites were enriched in 3’-UTR as well as 5’-UTR. Similar enrichment patterns were observed in TDP-43-CLIP sites (data from Hallegger et al. 2021). TDP-43-ADAR showed slightly lower 3’-UTR enrichment, however if the sites were shifted by 50bp there was a 16% increase in the number of sites in the 3’-UTR.

CLIP experiments indicate that TDP-43 binds to introns as well as to the 3’-UTRs of target transcripts (Polymenidou et al. 2011; Hallegger et al. 2021). Since the Smart-seq libraries generated in this study are generated from poly-A-mRNA, introns should and do represent a very small portion of sequenced transcripts. Editing sites were however enriched in the 3’-UTRs of target transcripts. This was shown by mapping TDP-43-ADAR or TDP-43-APOBEC editing sites as well as TDP-43 CLIP binding sites (Hallegger et al. 2021) to the genome and calculating the percentage of sites within the 5’-UTR, coding sequence and 3’-UTR. This distribution was compared to the transcriptome distribution and graphed as fold change relative to that distribution (Fig. 3C).

All three samples show strong enrichment in 5’-UTRs as well as 3’-UTRs with an underrepresentation of sites in the coding sequence (cds). A higher percentage of TDP-43-ADAR editing sites map to the cds than those from TDP-43-APOBEC and TDP-43-CLIP. However, if the coordinates of the TDP-43-ADAR editing site are expanded by 50bp, 16% of the TDP-43-ADAR editing sites shift from being the cds to the 3’-UTR (Fig. 3C; right). This suggests that many of these binding events occur in the 3’UTR near the cds-3’-UTR border.

The 3’-UTR and 5’-UTR enrichment of TDP-43-ADAR, TDP-43-APOBEC and TDP-43-CLIP sites are quite similar. To determine whether these three methods identify the same binding regions, we expanded each editing or CLIP site by 100bp in both directions and then compared the three regions (Fig. 4A and B). 47% of the TDP-43-APOBEC and 30% of the TDP-43-ADAR sites are in the same genes and overlapping (Fig. 4A; blue), whereas the other edited regions were unique to TDP-43-APOBEC and TDP-43-ADAR (Fig. 4A; green). We then asked whether editing sites that are present in both TDP-43-ADAR and TDP-43-APOBEC are more likely to also be identified by CLIP. Indeed, these common sites are 3-4-fold increased in the percentage that are also identified in CLIP experiments (blue; Fig. 4B).

**Figure 4.**
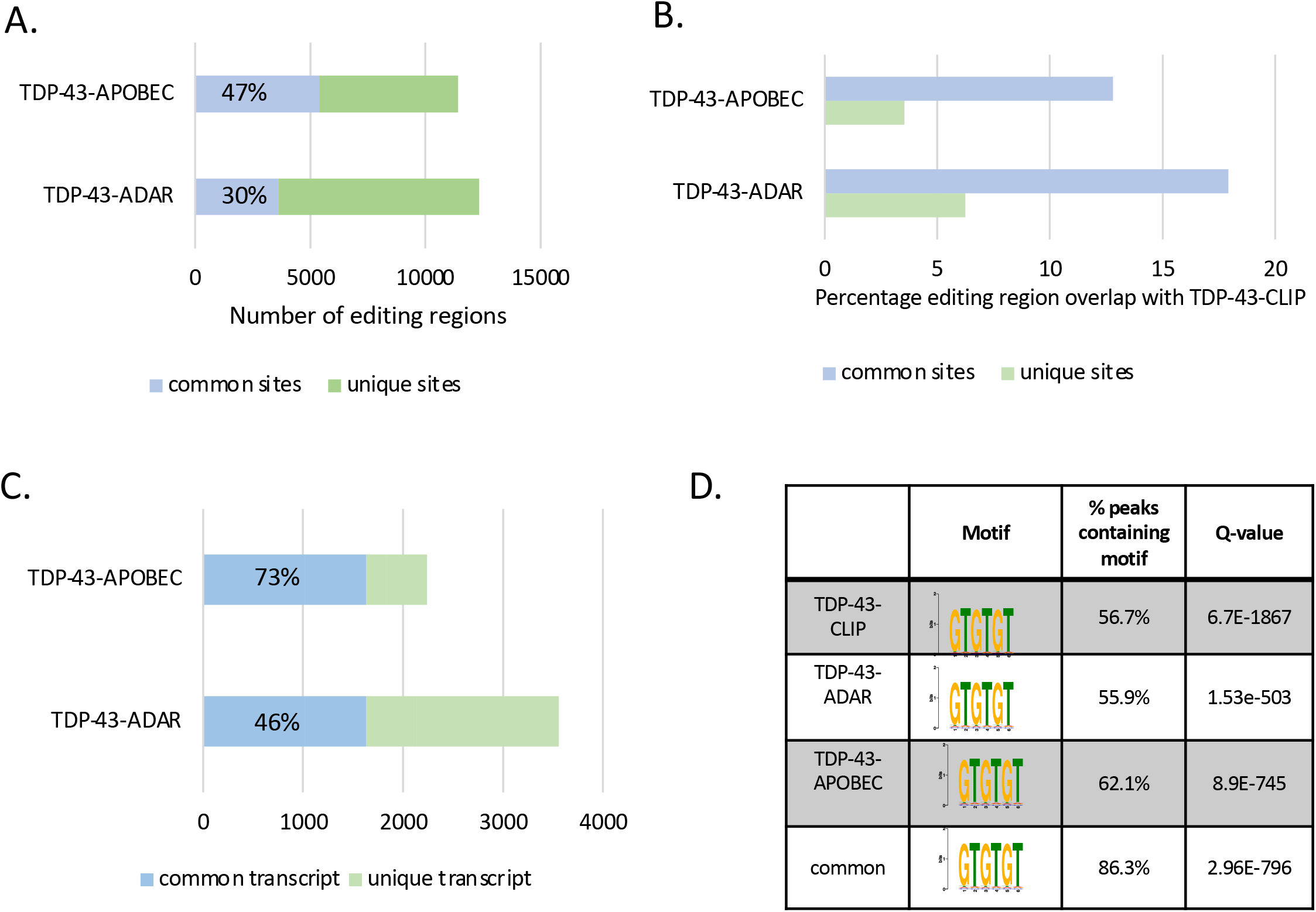
TDP-43-ADAR and TDP-43-APOBEC identified common edited regions and transcripts. A. TDP-43-ADAR and TDP-43-APOBEC identify common regions. The coordinates of identified editing sites were expanded by 50bp in each direction and examined for overlapping regions. Between 30-47% of the edited regions found in TDP-43 TRIBE and TDP-43 STAMP were found using both methods (blue). Those editing regions that were not found in common and were unique to either TDP-43-ADAR or TDP-43-APOBEC are show in green. B. Regions edited by both TDP-43-ADAR and TDP-43-APOBEC were more likely to be identified as a RBP binding site using CLIP. Between 10-20% of the 100bp regions containing editing sites generated by TDP-43-ADAR and TDP-43-APOBEC are also identified in TDP-43-CLIP experiments (blue). C. Many transcripts were identified as targets of TDP-43 using both TDP-43 TRIBE and TDP-43 STAMP. 70% of the transcripts identified by TDP-43-APOBEC were also identified by TDP-43-ADAR (blue). 46% of those transcripts identified by TDP-43-ADAR were also identified by TDP-43-APOBEC. D. Regions identified as edited by both TDP-43-ADAR and TDP-43-APOBEC are enriched in GU/GT-rich motifs favored by TDP-43 for binding. The 100bp region surrounding RNA editing sites generated by TDP-43-ADAR, TDP-43-APOBEC, and TDP-43-CLIP sites was analyzed using XStreme (Grant and Bailey 2021). 50-60% of the regions edited by TDP-43-ADAR, TDP-43-APOBEC as well as those containing TDP-43-CLIP signals contain GU/GT-rich motifs with high significance. Those edited regions that were identified by both TDP-43-ADAR and TDP-43-APOBEC showed a higher inclusion of the GU/GT-rich motif; 86% of the peaks contain this motif.

Because RBP-ADAR fusions often edit nucleotides as far as 500 nt from the RBP binding site (Xu et al. 2018), TDP-43-ADAR and TDP-43-APOBEC may identify similar transcripts even if their editing sites are not within 200bp of one another. We therefore compared the three target transcript lists: 73% of the TDP-43-APOBEC transcripts and 46% of the TDP-43-ADAR transcripts are identified by both methods (Fig. 4C; blue).

Because TDP-43-ADAR and TDP-43-APOBEC identify a substantially overlapping set of transcripts, the two methods can probably be combined to identify a higher confidence set of TDP-43 targets as also suggested by the increase overlap between CLIP and the common edited regions. We therefore examined the 100bp regions surrounding TDP-43-ADAR, TDP-43-APOBEC and TDP-43-CLIP sites for the GU-rich motif that is associated with TDP-43-binding (Polymenidou et al. 2011; Tollervey et al. 2011). Since the GUGUGU or GTGTGT motif was originally identified in CLIP experiments, we first examined the 100bp region surrounding TDP-43-CLIP sites. 57% of these sites contain a GTGTGT motif, a highly significant enrichment (Fig. 4D; q=6.7E-1867). The TDP-43-ADAR and TDP-43-APOBEC identified regions were also significantly enriched for GTGTGT motifs; a similar percentage of these regions, 56% and 62%, contain GTGTGT motifs (Fig. 4D). For regions identified by both methods, the GTGTGT motif was identified in 86% of the binding regions with even higher statistical significance (Fig. 4D). This suggests that higher confidence binding regions are indeed identified by assaying both fusion proteins rather than just one.

### Comparison of TRIBE and STAMP in *Drosophila*

We decided to follow the success of combining the ADAR and APOBEC editing enzymes to identify higher confidence TDP-43 targets in mammalian cells by testing the two enzymes side-by-side in *Drosophila* Schneider 2 cells (*Drosophila* S2 cells). To this end, we adopted the same strategy used above in mammalian cells and replaced the ADARcd with the rat APOBEC1 coding sequence. TRIBE was originally developed in *Drosophila*, and we assayed the same two RBPs previously used to identify TRIBE targets in S2 cells, namely, Hrp48 and Thor (*Drosophila* EIF4E-BP; McMahon et al. 2016; Jin et al. 2020). This resulted in a panel of plasmids that contained the Metallothionein inducible promoter (MT) driving a RBP-editing enzyme fusion followed by a p2A self-cleaving peptide and dsRed for visualization (Fig. 5A). We also generated enzyme only and dsRed only controls using the same general strategy. We transfected these expression plasmids into *Drosophila* S2 cells and isolated dsRed-positive cells via FACS (BD Melody). Poly-A mRNA was isolated from 400 cells and was used as input for RNA sequencing libraries generated using Smart-seq2 (Picelli et al. 2014). RNA editing sites were identified using the new TRIBE computational pipeline described above adapted for *Drosophila* (see Materials and Methods).

**Figure 5.**
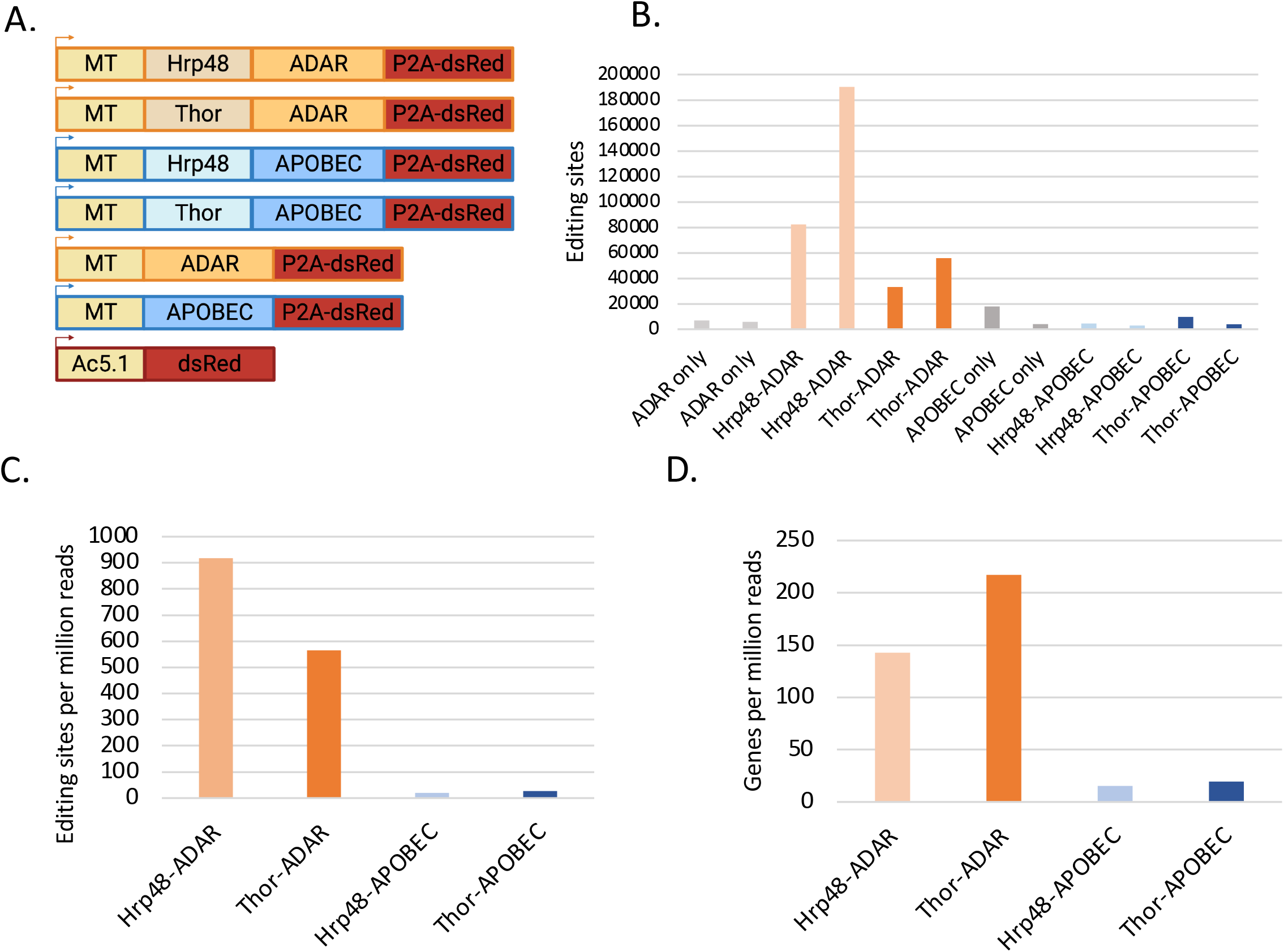
STAMP does not work well in *Drosophila* S2 cells. A. Schematic of the expression constructs used to test TRIBE and STAMP in *Drosophila* S2 cells. B. Number of editing sites identified using Hrp48 and Thor TRIBE (orange) and STAMP (blue) in two biological replicates. ADAR and APOBEC only controls are also shown to illustrate background editing of each enzyme (gray). Hrp48-ADAR and Thor-ADAR generated ~8-fold more editing sites than the ADAR-only control. In contrast, Hrp48-APOBEC and Thor-APOBEC did not show an enrichment of editing sites relative to the APOBEC only controls. C. Hrp48-APOBEC and Thor-APOBEC generate very few consistent editing sites. Editing sites identified in both biological replicates and not found in enzyme only controls were quantified and normalized relative to the total number of reads in the RNA sequencing library. D. Hrp48 and Thor STAMP identified very few target transcripts in *Drosophila* S2 cells compared to Hrp48 and Thor TRIBE.

As previously shown, expression of both Hrp48-ADAR and Thor-ADAR dramatically increased the number of editing sites above the low level of editing achieved by expressing the ADARcd alone (Fig. 5B; orange and light gray; McMahon et al. 2016; Jin et al. 2020). In contrast and unlike in human cells (Fig. 1B), expression of Hrp48-APOBEC and Thor-APOBEC did not cause an increase in editing above enzyme-only levels (Fig. 4B; blue and dark grey). This was not due to the instability of the APOBEC fusion proteins; western blotting could easily detect Hrp48-APOBEC (Sup. Fig.2). Moreover, the few C to T transitions appear to be due to APOBEC; these editing events are much more prevalent than other nucleotide changes and show the characteristic site choice of this enzyme (flanking A or T nucleotides; Sup. Fig. 3). This suggests that both RBP-APOBEC fusions only inefficiently edit target mRNAs in *Drosophila* tissue culture cells. Indeed, the number of RBP-APOBEC editing sites and mRNA targets that passed threshold (consistent between the two biological replicates and not edited at greater than 1% editing in the enzyme-only controls) was more than 10-fold less than the same RBPs fused to ADAR (Fig. 5C and D).

The experiments in human cells shown above indicated that RBP-APOBEC fusions were likely to edit more sites but with a lower percentage editing (Fig. 2C). This same trend is observed in *Drosophila*; the mean editing percentage on Hrp48-APOBEC editing sites is 8.8% while the mean editing of Hrp48-ADAR sites is significantly higher (26.1%). To test the possibility that requiring 6% editing was preventing the identification of RBP-APOBEC editing sites, we re-analyzed the data with a 4% editing cutoff (Sup. Fig 4). Although the number of editing sites identified by Thor-APOBEC and Hrp48-APOBEC increased approximately 3-fold, the number of APOBEC-only editing sites increased 15-fold. This suggests that a too high editing threshold is not the reason for the weak RBP-APOBEC editing relative to APOBEC-only editing. We do not know why RBP-APOBEC chimeric proteins do not work well in *Drosophila*; it may be due to the absence of the important mammalian APOBEC cofactors AICF and RBM7 or to temperature differences (23°C versus 37°C) (reviewed in Pecori et al. 2022; see Discussion).

## Discussion

RNA binding proteins guide RNAs throughout their life, by binding to nascent RNAs as they emerge from the polymerase, by facilitating the removal of introns and nuclear export, and then by modulating mRNA turnover, localization, and translation in the cytoplasm. Mutations in over 1000 RBPs are linked to human diseases including Fragile X and ALS (reviewed in (Gebauer et al. 2021). Therefore it is critical to have reliable tools to identify the mRNA targets of RBPs, in specific cell types and even single cells. To this end, we directly compared in this manuscript TRIBE (RBP-ADAR) and STAMP (RBP-APOBEC) in both human cells and in *Drosophila* cells. In human cells, TDP-43 targets were successfully identified using both methods, and approximately 70% of the STAMP targets were identified by TRIBE. The results also indicated that a higher confidence set of RBP targets is identified by defining the common targets identified with both methods.

Consistent with our finding that TRIBE and STAMP both work well and similarly is a very recent study that utilized both TRIBE and STAMP simultaneously, which the authors dubbed TRIBE-STAMP (Flamand et al. 2022). In this study Flamand et al., investigated the mRNA targets of the m^6^A reader proteins, YTHDF1, YTHDF2 and YTHDF3 and showed that these three proteins identified similar target mRNAs with both TRIBE and STAMP. Although the overlap between these two methods was greater than observed here, this may be because the m^6^A reader proteins bound a very large percentage of the transcriptome, perhaps generating a higher likelihood of common targets. These authors also showed that TRIBE and STAMP can be combined to determine if two RPBs bind to the same transcript, another application of using both TRIBE and STAMP in parallel.

Although we were hopeful STAMP would also become a second tool for use in *Drosophila*, we were unable to observe significant RNA editing by APOBEC fused to two different RBPs in *Drosophila* S2 cells. The *Drosophila* constructs mirror those successfully used in human cells and in the original STAMP study, i.e., APOBEC is fused to the C-terminus of the RBP. The STAMP fusion proteins are transcribed and translated, and some APOBEC-driven editing is detected (Fig. 5B and Sup. Fig. 3). However, the editing is quite poor; the Thor-APOBEC and Hrp48-APOBEC editing levels are similar to the APOBEC-only controls (Fig. 5B).

A recent publication on BioRxiv suggests a possible explanation for this lack of APOBEC editing activity in S2 cells. In this manuscript, the authors fused APOBEC1 to Cas9 in an attempt to use APOBEC as a gene editing enzyme in *Drosophila*. APOBEC deaminase activity was poor at lower temperatures (18°-24°) but functional at 29°C (Doll et al. 2022). This may be the explanation for low APOBEC deaminase in *Drosophila* S2 cells cultured at 23°C. It further suggests that STAMP may not be a viable option for detecting RBP target mRNAs in other organisms or systems requiring lower growth temperatures.

Does either TRIBE or STAMP pose a significant advantage for RBP target identification in mammalian cells? Does one method identify substantially more false negatives or false positives than the other?

As far as false negatives are concerned, each enzyme has intrinsic features that bias the editing of an RBP-bound RNA. TDP-43-APOBEC has a strong nearest neighbor preference as nearly all cytosines edited are flanked by A or T (84%; Fig. 2F). A previous study has argued that APOBEC is ideal for RBP target mRNA identification as it can edit cytosines in single stranded mRNA which should be between 25-35% of nucleotides in mammalian transcripts (Brannan et al. 2020). However, the strong nearest neighbor preference observed both here and in a previous study (Rosenberg et al. 2011) suggests that RBP-APOBEC fusion proteins edit a more limited set of cytosines. Nonetheless, the observation that TDP-43-APOBEC tends to edit different cytosines on the same transcript or multiple copies of the same transcript suggests that this nearest neighbor preference is not an issue for the identification of most mRNA targets and target regions.

In contrast, TDP-43-ADARcd is more likely to edit the same adenosine resulting in the editing of the exact same nucleotide in multiple experiments (Fig. 2A). Since the ADARcd has less stringent nearest neighbor preferences than APOBEC (Fig. 2E and 2F), the repeat editing of specific adenosines by the ADARcd is more likely due to its double-stranded RNA requirement (Macbeth et al. 2005) and may be responsible for the fewer TRIBE edits compared to STAMP edits: an average of 4.8 sites/transcript for the ADARcd versus 7 sites/transcript for APOBEC (Fig. 2D). Despite this modest difference, TRIBE identifies somewhat more target transcripts than STAMP (Fig. 3B). This indicates that the ADARcd double stranded RNA requirement is not an obstacle to target identification as previously discussed (McMahon et al. 2016) and that TRIBE does not suffer from a false negative problem relative to STAMP.

A previous study has purported that TRIBE is not well suited for RBP target identification compared to STAMP because the ADARcd only generates very few edits, only 1 to 2 per target transcript, and is incapable of editing coding regions (cds) due to their single stranded nature (Brannan et al. 2020). This comment was based on results with Fmr1 using the first version of TRIBE prior to the adoption in 2018 of an ADARcd containing the E488Q mutation (Xu et al. 2018). This HyperTRIBE method gives rise to much more editing, due to faster editing speed and less preference for an UAG neighboring sequence surrounding the editing site (Kuttan and Bass 2012). In fact, as shown here and elsewhere, TRIBE identifies target transcripts with editing sites ranging from 1 to 43 nucleotides and also edits cds (Fig. 2D and Cheng et al. 2021).

What about false positives, editing sites that do not reflect RBP mRNA binding? TRIBE only uses the ADAR catalytic domain (ADARcd), avoiding its RNA binding regions. As a consequence, the ADARcd alone generates a low level of A to G editing, suggesting that most TDP-43-ADARcd editing sites and target transcripts are real positives. In contrast, the catalytic and RNA binding regions of APOBEC are not defined, requiring use of the entire protein in STAMP. This likely explains the 5-fold increase in editing by APOBEC-only compared to ADAR-only (Fig. 1B). APOBEC may normally bind to and edit targets with the help of two endogenous cofactors, AICF and RBM7 (reviewed in (Pecori et al. 2022). It is therefore possible that APOBEC and its cofactors compete with TDP-43 for binding site recognition.

The ability of APOBEC to edit single stranded DNA could also impact the level of detectable background editing. Indeed, rat APOBEC1 used here and in the original STAMP study has also been used successfully in combination with CRISPR for genome editing (Komor et al. 2016; Komor et al. 2017). Other studies have shown that APOBEC1 can drive off-target DNA editing and RNA-editing even when fused to Cas9 (Grunewald et al. 2019; Jin et al. 2019; Kanca et al. 2019; McGrath et al. 2019). This observation was true even with transient transfection of the Cas9-Apobec fusions (McGrath et al. 2019). To date, genomic DNA has not been examined in STAMP, making it uncertain whether APOBEC-driven single stranded DNA editing is contributing to false positive editing sites and transcripts. Although current computational approaches cannot distinguish DNA-editing from RNA editing, we attempted to ameliorate this issue by eliminating from the final list of editing sites any nucleotide that has greater than 1% editing in enzyme-only control samples.

Other possible sources of false positive editing sites and target transcripts are shared by TRIBE and STAMP. First, these methods currently rely on overexpression, which can cause RBPs to bind to and edit secondary, low efficiency targets as well as primary high efficiency targets. Second, there are false-positive sources of editing that are intrinsic to living cells, endogenous editing by ADAR and APOBEC as well as genomic variation in the form of sNPs. Unlike computational approaches that identify editing sites by comparing the experimental RNA sample to publicly available genomic sequences however, our computational pipeline is set up to handle these issues and is comprehensive for STAMP as well as TRIBE. By directly comparing RNA sequences from TRIBE and STAMP samples to the same cell line expressing only GFP, we were able to eliminate most of these false positive sources, i.e., sites that are edited by endogenous ADAR and APOBEC as well as sNPs are present in the control cell line (see Materials and Methods). A third source of false positives is technical error, whether from PCR during RNA sequencing library generation or from sequencing itself. To control for this, we now require edited nucleotides to have 20 reads of coverage and greater than 6% editing to ensure that a candidate altered nucleotide is genuine. Although this requirement requires additional sequencing depth (at least 12 million uniquely mapped reads for each sample) to detect editing events in less abundant transcripts, it prevents the identification of an editing site due to a single A to G (ADAR) or C to T (APOBEC) change that could be due to technical error. Although a 6% editing cutoff does reduce the number of editing sites and target transcripts identified, reducing this cutoff to 5% indicates that 6% is not too high: The lower cutoff caused an increase in all nucleotide transitions, not specifically A to G (ADAR) or C to T (APOBEC). More generally, our new, comprehensive pipeline is conservative and designed to reduce the contribution of false positives. This also means that the number of bona fide editing sites and transcripts could be considerably greater than what we define here.

The topics of false positives and false negatives warrants a return to considering the lack of overlap between TRIBE, STAMP and CLIP (Fig. 4). There is unfortunately no ground truth as CLIP itself is prone to false positives and false negatives (reviewed in Xu et al. 2022). It is therefore difficult to ascertain whether one of these methods is more accurate than another. In light of this background, a conservative approach is to identify a set of high confidence targets by overlapping methods. Approximately 1600 transcripts were identified by both TDP-43-ADAR and TDP-43-APOBEC (Fig. 4C). Although it is difficult to evaluate which of these targets are most relevant, it is notable that edited regions identified by both TRIBE and STAMP have a higher frequency of the GTGTGT binding motif than TDP-43-ADAR, TDP-43-APOBEC or TDP-43-CLIP alone (Fig. 4D). They are also more likely to be identified as TDP-43-CLIP targets (Fig. 4B). In addition, an irrefutable TDP-43 target, the TDP-43-mRNA itself, is found in both TRIBE and STAMP. Although other known neuronal targets of TDP-43 are not expressed in HEK293T cells, two S2 cell targets are also identified by both TRIBE and STAMP (Rac1, futsch/map1b). We therefore suggest that a good approach in the future to identify high confidence RBP targets in mammalian systems is to examine targets using both TRIBE and STAMP and move forward with the common targets. This will be especially useful when CLIP is not easy to apply.

## MATERIALS AND METHODS

### Plasmids

The details of human and *Drosophila* plasmid construction are below and listed in Supplemental Table 1. For all plasmids, we used either Q5 (NEB) or Extaq (Takara) DNA polymerases to PCR amplify inserts. All inserts and vectors were gel purified using the gel extraction kit (QIAGEN). We inserted all fragments into vectors using either Gibson Assembly (NEB) or NEBuilder (NEB) unless otherwise noted. All plasmids were transformed into NEBalpha DH5 high efficiency competent cells. We verified candidate plasmids using colony PCR using primers listed in Supplemental Table 2 and REDTaq ReadyMix (Sigma-Aldrich). All plasmids were miniprepped (QIAGEN) and sequenced with Plamidsaurus (https://www.plasmidsaurus.com).

#### HEK-293 expression vectors

Plasmids for expressing TDP-43-ADAR (pCMV-hADARcd-E488Q) as well as the ADAR only control (pCMV-hADARcd-E488Q) were previously published (Herzog et al. 2020). To generate pCMV-APOBEC (pCR25), we amplified rat APOBEC1 from pMT-Hrp48-APOBEC-P2A-dsRed (pCR8; see below) with primers CR60 and CR61 and inserted it into pCMV-ADARcd-E488Q digested with EcoRI and KpnI. To clone, pCMV-TDP-43-APOBEC (pCR27) we amplified APOBEC from pMT-Hrp48-APOBEC-P2A-dsRed (pCR8) with primers CR73 and CR74 and inserted into pCMV-TDP-43-ADARcd-E488Q linearized using PCR with primers CR70 and CR71.

#### *Drosophila* plasmids

A *Drosophila* HyperTRIBE plasmid was generated by including a selfcleaving peptide, p2A, followed by dsRed downstream of the Hrp48-ADAR sequence (pMT-Hrp48-ADAR-E488Q-P2A-dsRed; pCR1). This plasmid and all related plasmids have a V5-tag 5’-of the editing enzyme. The plasmid pMT-Tyf-ADAR-E488Q-P2A-dsRed was digested with PmeI and NotI to liberate ADAR-E488Q-P2A-dsRed. This fragment was ligated into PmeI and Not1-digested pMT-Hrp48-ADAR-E488Q using T4 ligase (NEB). pMT-Hrp48-APOBEC-P2A-dsRed (pCR8) was made by amplifying rat APOBEC1 from pCMV-BE1 (Addgene #73019) using primers CR26 and CR27. Gibson Assembly was then used to insert APOBEC1 into pMT-Hrp48-ADAR-E488Q-P2A-dsRED (pCR1) digested with ApaI and NotI to remove ADAR-E488Q. The resulting plasmid was then cut with NotI to insert a linker (made by annealing primers CR29 and CR30) using Gibson assembly. To clone the APOBEC-only control (pMT-APOBEC-p2A-dsRed; pCR10), we amplified APOBEC from pCMV-BE1 (Addgene #73019) using primers CR28 and CR27. Plasmid pMT-Hrp48-ADAR-P2A-dsRed (pCR1) was digested with ApaI and KpnI to liberate Hrp48-ADAR and APOBEC was inserted using Gibson Assembly. To clone an ADAR-only control (pMT-ADAR-P2A-dsRed; pCR12), we digested pMT_Hrp48_ADAR_E488Q_P2A_dsRed (pCR1) with NotI and KpnI to remove Hrp48. We blunted the resulting sticky ends using Klenow end-blunting (NEB) and re-circularized the plasmid using T4 blunt-end ligation (NEB). To clone pMT-Thor-APOBEC-P2A-dsRed (pCR26) we amplified Thor from pMT-Thor-Linker-HyperTRIBE (Jin et al. 2020) with primers CR66 and CR67 and inserted it into pMT-Hrp48-Linker-APOBEC-P2A-dsRed (pCR8) digested with KpnI and NotI to remove Hrp48 using NEBuilder (NEB). To clone pMT-Thor-ADAR-P2A-dsRed (pCR30) we amplified Thor from pMT-Thor-Linker-HyperTRIBE (Jin et al. 2020) using primers CR80 and CR81 and inserted into pMT-Hrp48-ADAR-E488Q-P2A-dsRed (pCR1) digested with KpnI and NotI to remove Hrp48. Insertion was done using NEBuilder.

### Cell Culture

HEK293T cells (ATCC CRL-3216) were cultured at 37°C in Gibco DMEM, high glucose, GlutaMAX supplement, pyruvate (Thermofisher) with 10% Fetalgro synthetic FBS (RMBio, FGR-BBT) and 1% Penicillin-Streptomycin (Genesee Scientific). HEK cells were transiently cotransfected with pCMV-EGFP and the relevant plasmid of interest in a six well plate using the Lipofectamine3000 protocol (Thermofisher). The CMV promoter is constitutively expressed and 24hrs after transfection, GFP-positive cells were collected with BD FACS Melody.

*Drosophila* S2 cells were cultured at 23°C in Schneider’s media with 10% Gibco HI-FBS (Thermofisher) and 1% Gibco Antibiotic-Antimycotic (Thermofisher). *Drosophila* S2 cells were transiently transfected using the Mirus TransIT-2020 transfection protocol (Mirusbio) in a 6 well plate for approximately 24 hours. Cells were induced for 24hrs by inducing the metallothionein promoter (pMT) using 0.5mM Copper sulfate (Sigma). DsRed-positive cells were collected with the BD FACS Melody.

### RNA Library Preparation

400 cells were sorted directly into aliquots of 100uL lysis buffer (Invitrogen Dynabeads mRNA Direct Kit;Thermofisher). Poly-A-plus RNA was isolated using Invitrogen Dynabeads mRNA Direct Kit (Thermofisher, 61012) and sequencing libraries were prepared following SmartSeq2 optimized protocol (Picelli et al. 2014). cDNA was quantified with D5000HS Tapestation (Agilent) and final tagmented libraries were quantified on D1000HS tapestation (Agilent). Libraries were sequenced on Illumina NextSeq500 with 75cycle high Output kit v2.5.

### TRIBE Pipeline

To identify editing sites, the TRIBE pipeline was used as previously described (Rahman et al. 2018); https://github.com/rosbashlab/HyperTRIBE). Briefly custom scripts were used to trim (Cutadapt; Martin 2011) and align (STAR; Dobin et al. 2013) reads to the appropriate genome (Human (GRChg38.p13) or *Drosophila* (dm6)). Mapped reads were used to generate gene expression data (see below) as well as bigwig files for visualization on the Integrated Genomics Viewer (IGV; Robinson et al. 2011). The mapped reads were then converted in a matrix file listing the number of As, Gs, Cs, and Ts found in sequencing reads at each genomic coordinate. This data was loaded into a mysql database for efficient querying. Editing sites were identified by identifying coordinates in the control samples (EGFP control for HEK-293 cells and the dsRed control for *Drosophila* S2 cells) that meet three criteria: 1) at least 80% of the reads were a A (ADAR) or a C (APOBEC), 2) less then 0.5% of the reads at that location were the edited base i.e., a G (ADAR) or a T (APOBEC), and 3) there were at least 9 reads covering the location. If these three conditions were met, then these coordinates were examined in the experimental samples. The coordinate is considered an editing site if: 1) there are at least 20 reads in the experimental sample, and 2) greater than 6% of the reads have an edited base (G (ADAR) and T (APOBEC)) at that location (see Rahman et al., 2018 and custom scripts). Supplemental Table 3 illustrates the effect of each filter on the total number of editing sites identified. By comparing experimental samples to a control RNA sample from the same cell line rather than publicly available genomic sequences, the majority of sNPs and endogenous editing events are filtered out in the initial editing site identification step.

Once editing sites were identified, a final list of editing sites for each TRIBE-RBP and STAMP-RBP was generated by identifying editing sites consistent between two biological replicates for RBP-ADAR and within 100bp of each other in two biological replicates RBP-APOBEC. Both approaches utilized bedtools intersect and bedtools closest(Quinlan 2014). Editing sites with at least 1% editing in the editing enzyme only control samples were identified using the custom script (Threshold_editsites_20reads.py) with editing threshold set to 0.01. The list of all sites identified with greater than 1% editing in either biological replicate of the enzyme only control libraries was then subtracted from the list of putative editing sites identified in the experimental samples using bedtools.

The number of editing sites per transcript was determined using a custom script (summarize_results.pl) which generates a list of all transcripts, the number of editing sites in that transcript (Rahman et al. 2018). Quantification of the distribution of RNA sequencing reads in the human and *Drosophila* RNA libraries was performed with read_distribution.py script in RSeQC4.0.0 (Wang et al. 2012). The distribution of editing sites between the 5’UTR, coding sequence (cds) and 3’UTR was determined using bedtools and custom scripts. Near neighbor preference was determined using custom scripts. To examine the overlap of TDP-43-TRIBE, TDP-43-STAMP and TDP-43-CLIP binding regions, the editing or CLIP site was expanded by 100 bp in both directions using bedtools slop. Then the overlap between the different regions was determined using bedtools intersect. All custom scripts generated for this study are annotated and deposited at https://github.com/rosbashlab/Comparison-of-TRIBE-and-STAMP.

To examine TDP-43-TRIBE, TDP-43-STAMP, and TDP-43-CLIP sites for the presence of GU-rich motifs preferentially bound by TDP-43, the region surrounding the editing site or CLIP site was expanded by 50bp using bedtools slop. The resulting bed file was used to retrieve sequence information for these regions to use as input for motif analysis using Xstreme (Grant and Bailey 2021).

### CLIP data analysis

CLIP data was downloaded from ArrayExpress with accession number E-MTAB-9436 (Hallegger et al. 2021). CLIP data from two biological replicates was analyzed using iCount at iMAPS (https://imaps.goodwright.com/history/). CLIP peaks identified in both biological replicates were identified using bedtools and used as a high-confidence set of putative TDP-43-CLIP sites.

### Gene Expression Analysis

Raw gene count values were generated using HTseq.scripts.count during initial mapping of RNA sequencing (Anders et al. 2015). Two replicates of either TDP-43-ADAR or TDP-43-APOBEC were compared to two replicates of the negative control cells (HEK-293 cells expressing only EGFP). Prior to differential expression analysis, data was filtered to include remove transcripts that were expressed at lower levels (expressed at less than 5 FPKM reads in more than 2 of the four samples). Differential gene expression upon overexpression of TRIBE and STAMP RBPs was analyzed using EdgeR (Robinson et al. 2010). The resulting smearplots are shown.

### Western Blot Analysis

To perform western blots, we adhered S2 cells transfected with pMT-Hrp48-APOBEC-P2A-dsRed (pCR8) and pMT-APOBEC-P2A-dsRED (pCR10) in wells of a 6 well plate and induced with 0.5mM copper sulfate for 24 hours. 250uL RIPA buffer (10mM HEPES pH7.5, 5mM Tris pH7.5, 50mM KCl, 10% glycerol, 2mM EDTA, 1% Triton X-100, 0.4% NP-40, protease cOmplete Mini tablet (Roche)) was added to each well, cells were scraped and lysed at 4°C for 15 minutes. After centrifugation (15,000rpm at 4°C for 10 minutes) samples were boiled in Laemmli buffer and loaded into a 10% MES NuPage Bis-Tris gel (Invitrogen) with the Novex™ SharpPre-stained Protein Standard (Invitrogen). The gel was transferred to nitrocellulose using iBLOT2 (Invitrogen) following manufacturer’s protocol. The blot was blocked with 5% milk in Tris-buffered saline with 1% Tween 20 (TBST) for 1hr before being incubated overnight at 4°C in and primary antibodies (1:1000 mouse anti-V5 Tag monoclonal Antibody (Invitrogen) and anti-Actin monoclonal antibody (Invitrogen) in 5% milk/TBST). Primary antibody was removed with 6 washes of TBST each for 5 minutes. The blot was incubated with secondary antibody (Cytiva Lifescience™ anti-Mouse IgG, peroxidase-linked species-specific whole antibody (from sheep); Fisher Scientific), diluted in 5% milk/TBST for one hour on a rocker at room temperature. The blot was then washed in 6x 5 minutes of TBST prior to being developed with Clarity™ Western ECL Substrate kit (Bio-Rad). To examine actin levels as a loading control blot was stripped using Restore™ WesternBlot Stripping Buffer (Thermo Scientific) for 15 minutes and re-blocked with 5% milk for 1hr at RT on rocker. Blots were imaged using a ChemiDoc (Biorad).

## DATA AVAILABILITY

All data generated in this study is deposited in Gene Expression Omnibus (GEO) under accession numbers_GSE223557. Human data is in subseries <u>GSE223555</u> and *Drosophila* data is in subseries <u>GSE223556</u>. The TRIBE analysis pipeline is publicly available at https://github.com/rosbashlab/HyperTRIBE). All scripts used in this manuscript are available at https://github.com/rosbashlab/Comparison-of-TRIBE-and-STAMP.

## ACKNOWLEDGEMENTS

We would like to thank all members of the Rosbash lab for helpful comments on this work. We thank Shlesha Richhariya and Don Rio for comments on the written manuscript. We thank Daniel Shin for lab management and administrative support. This work was supported by the Howard Hughes Medical Institute and the NIH R01 (DA037721).

## FIGURE LEGENDS

**Supplemental Figure 1.**
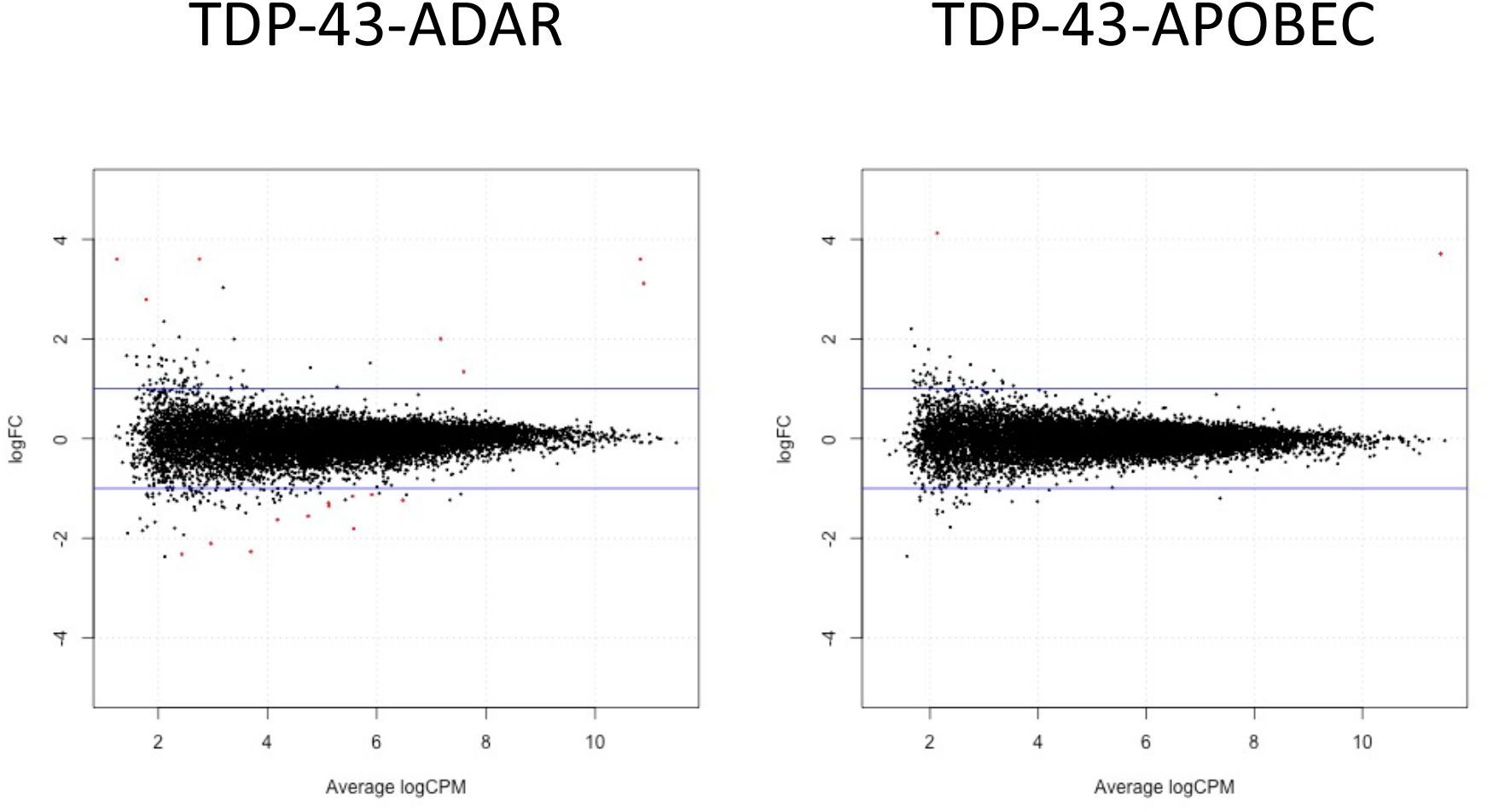
TDP-43-ADAR and TDP-43-APOBEC did not dramatically perturb HEK-293 cell transcriptomes. Smear plots generated by EdgeR analysis comparing HEK cells expressing GFP to HEK cells expressing TDP-43-ADAR (left) or TDP-43-APOBEC (right). All transcripts expressed at greater than 5 FPKM in at least two experiments included in the analysis. Gray lines indicate a threshold for a 2-fold change. Significantly altered genes (q< 0.05) are indicated as red dots. Very few genes changed with the expression of either transgene.

**Supplemental Fig.2.**
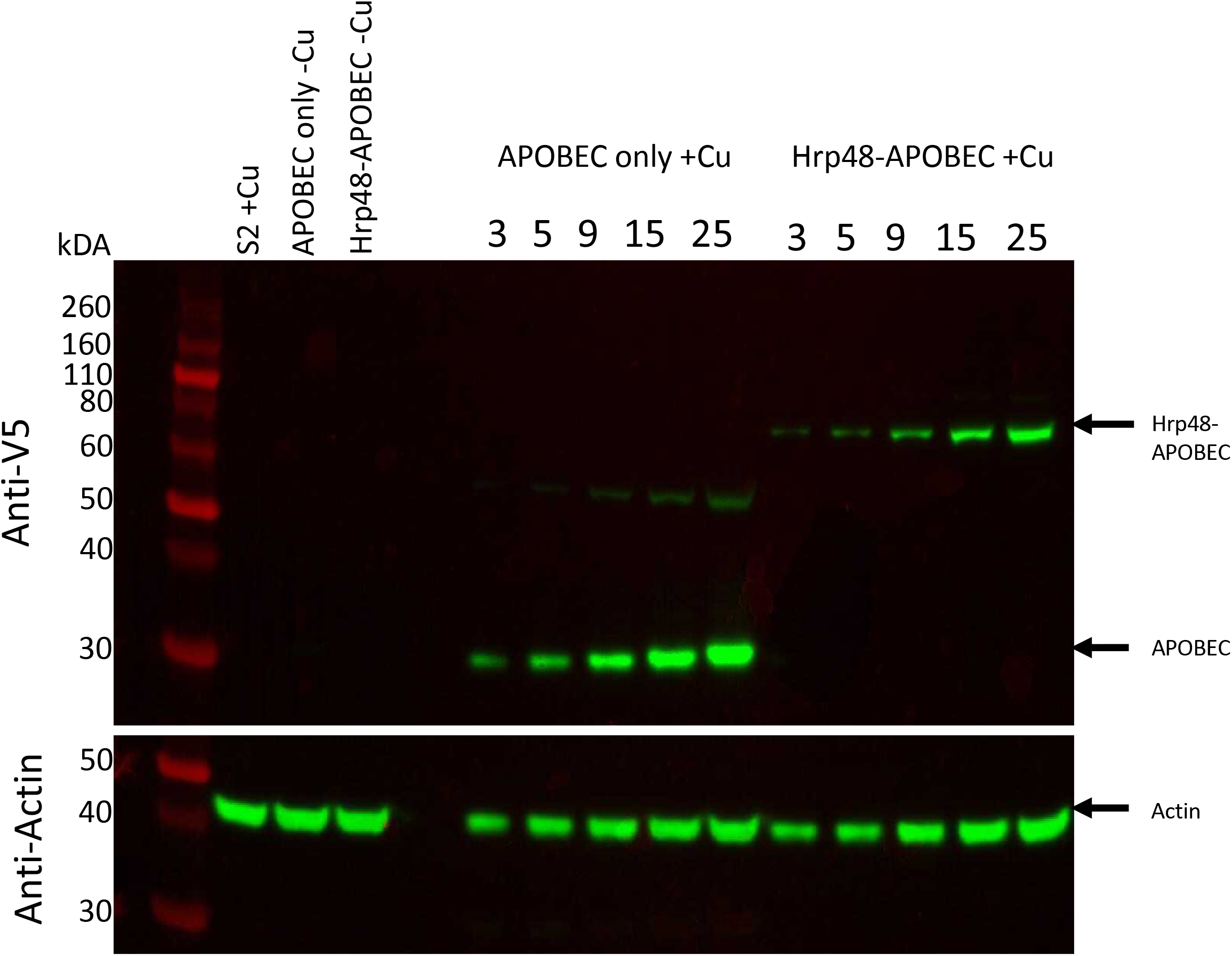
Hrp48-APOBEC and APOBEC proteins were expressed in *Drosophila* S2 cells. Anti-V5 (TOP) and anti-Actin (bottom) western blots of protein extracts from *Drosophila* S2 cells expressing either APOBEC only or Hrp48-APOBEC under the control of the copper inducible metallothionein promoter. Wild-type *Drosophila* S2 cells induced with copper are shown in lane1 as a negative control. Additional negative controls are shown in lane 2 and 3: APOBEC only and Hrp48-APOBEC transfected cells without copper induction. Lanes 5-9 show increasing amounts of copper induced APOBEC only cell lysates (3μl to 25μL). Lanes 11-15 show increasing amounts of copper induced Hrp48-APOBEC cell lysates (3μl to 25μL). APOBEC and HRP48-APOBEC were both tagged with a V5 tag and western blotting shows bands at the expected sizes. Anti-actin blot (bottom) included as a loading control.

**Supplemental Figure 3:**
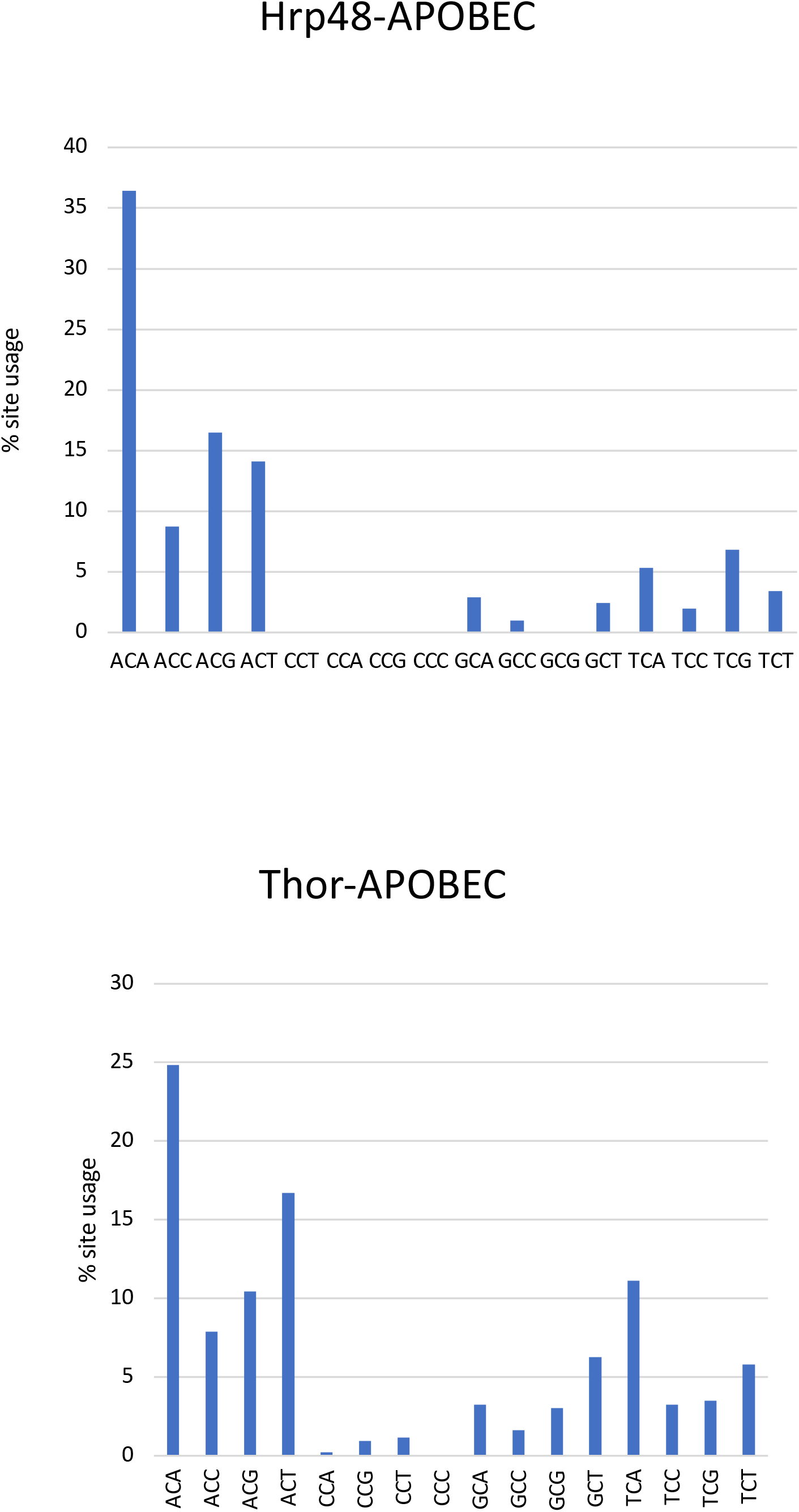
Editing sites generated by expression of Hrp48-APOBEC and Thor-APOBEC show bias toward cytosines flanked by A or T. However, this bias is not as strong as observed in mammalian cells.

**Supplemental Figure 4:**
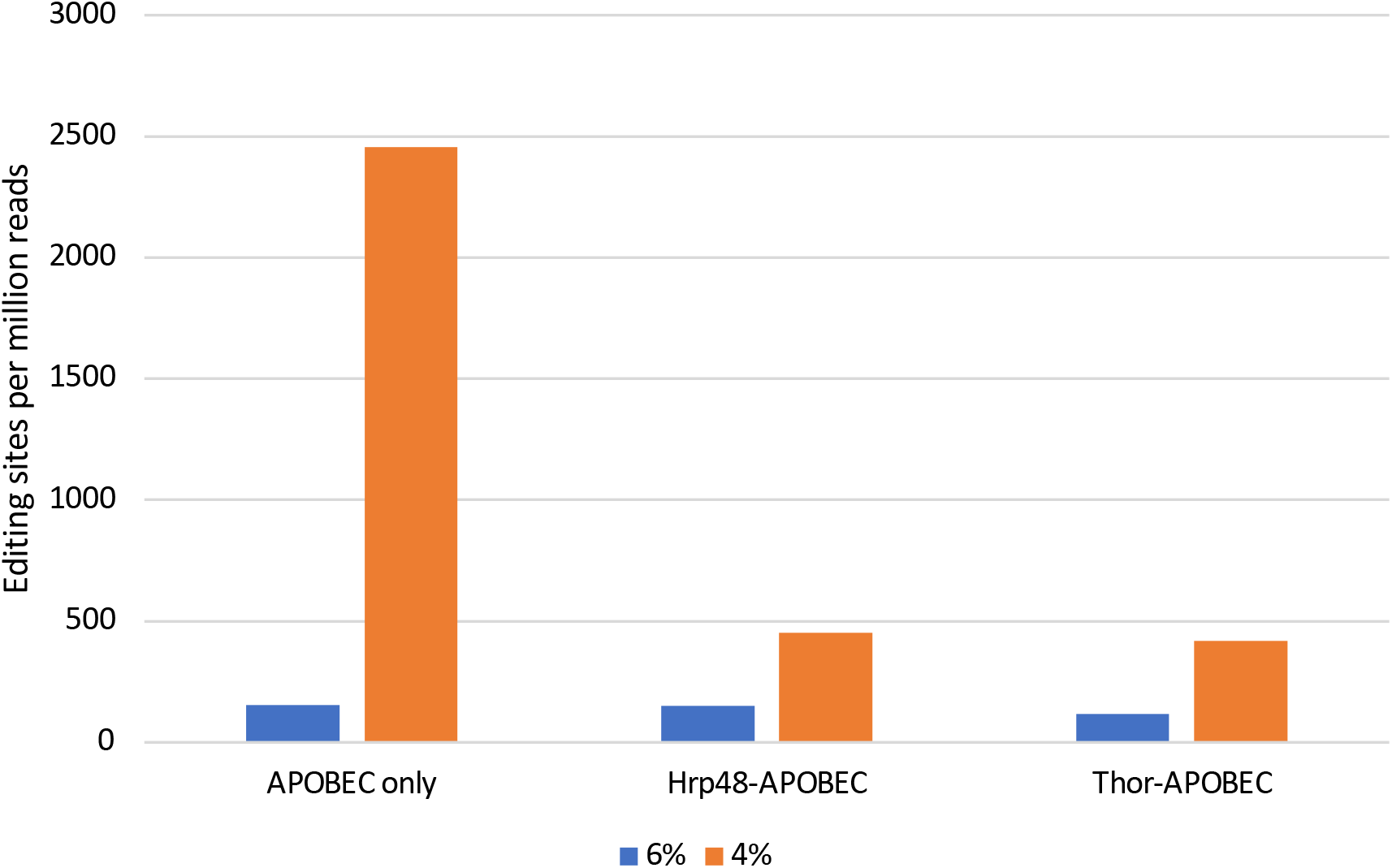
Reducing the editing threshold from 6% to 4% disproportionally affected the number of APOBEC only editing sites identified relative to Hrp48-APOBEC and Thor-APOBEC. Since RBP-APOBEC fusions showed lower percentages of editing at a particular cytosine nucleotide, we reduced the threshold for calling an editing site to four percent to see if this would allow for more RBP-APOBEC-driven editing sites in *Drosophila*. However, the number of editing sites found when APOBEC alone was overexpressed increases much more dramatically. This suggests that tethering APOBEC to a RBP in *Drosophila* may actually decrease its editing activity.

**Supplemental Table 1: Plasmids used in this study.**

**Supplemental Table 2: Primers used in this study**

**Supplemental Table 3: Editing site filtering.** Chart illustrating different steps in editing site identification and how many sites are lost at each filtering step.

**Supplemental Data 1: Editing sites and target transcripts identified by HEK-293 cells.** Excel worksheet with tables listing the final editing sites and target genes identified in TDP-43-TRIBE and STAMP.

**Supplemental Data 2: Editing sites and target transcripts identified in *Drosophila* S2 cells.** Excel worksheet with tables listing the final editing sites and target genes identified in Hrp48 and Thor TRIBE and STAMP.

## Notes

https://github.com/rosbashlab/Comparison-of-TRIBE-and-STAMP.git

https://github.com/rosbashlab/HyperTRIBE.git

https://www.ncbi.nlm.nih.gov/geo/query/acc.cgi?acc=GSE223557

### Competing Interest Statement

Michael Rosbash holds a patent on TRIBE approach (US11401514B2). The authors nor the institution have received any payments or services from a third party that could be perceived as influencing this work.

